# Causal Associations of Circulating Helicobacter pylori Antibodies with 5 Skin Disorders: A Mendelian Randomization Analysis

**DOI:** 10.1101/2024.08.10.607427

**Authors:** Ao He, Yao Ni, Na Qiao, Wenli Huang, Fengshan Gan, Zhuo Gong, Xuesong Yang, Xian Zhao

## Abstract

Helicobacter pylori (H. pylori) infection, a prevalent bacterial ailment affecting the stomach, has been linked to diverse gastrointestinal disorders. Nevertheless, the correlation between H. pylori infection and skin disorders has not been definitively established. We use Mendelian randomization analysis to explore causal relationships with seven pylori antibody levels and five skin diseases, such as rosacea, psoriasis, alopecia areata, pruritus and urticaria. The primary analytical method employed was inverse variance weighted (IVW), and a sensitivity analysis was executed to evaluate the robustness of the primary findings. The results revealed a positive causal relationship between H. pylori catalase antibody levels and AA (IVW: OR = 1.32, 95% CI = 1.07 to 1.61, P = 0.008), as well as a positive causal relationship between H. pylori UreA antibody levels and pruritus (IVW: OR = 1.17, 95% CI = 1.05 to 1.31, P = 0.006). These effects were robust to sensitivity analyses. Our results suggested that H. pylori infection was a risk factor for AA and pruritus. This finding may have implications for the prevention and treatment of these skin disorders.

## 1. Introduction

Helicobacter pylori (H. pylori) is a spiral-shaped Gram-negative bacterium that infects more than 50% of the world’s population. It acts as a causative factor in certain gastrointestinal diseases, such as peptic ulcers and gastric cancer. Furthermore, it contributes to the etiology of various disorders beyond the gastrointestinal tract.[1,2] Such as iron deficiency anemia, idiopathic thrombocytopenic purpura, as well as chronic urticaria, and other skin diseases. H. pylori infection may lead to the occurrence of these diseases including systemic inflammatory responses, and stimulating autoimmunity. Several H. pylori antibodies have been identified in patients with H. pylori infection.

Research has demonstrated a correlation between elevated serum levels of H. pylori IgG and cytotoxin-associated gene-A (CagA) antibodies and an augmented susceptibility to specific skin disorders.[3,4] The efficacy of H. pylori eradication therapy has been observed in certain cases of skin disorders, including chronic urticaria, rosacea, psoriasis, and alopecia areata (AA).[5,6] Nevertheless, it is crucial to acknowledge that many of these investigations rely on case reports or trial studies, potentially leading to conflicting outcomes. Consequently, the association between H. pylori infection and skin disorders remains contentious within the current body of research.[7,8] The results of these studies are affected by sample size limitations, unavoidable confounding factors, and reverse causality, it is difficult for us to determine the potential causal relationship between H. pylori infection and related skin disorders.[9-11]

Mendelian randomization (MR) is a widely used epidemiological investigation method, utilizing genetic variation as an instrumental variable to deduce the causal association between exposure and outcome.[9] The core principle randomly assigning genetic variation to offspring at conception, ensuring that it is not affected by potential confounding factors and minimizing the impact of reverse causation. MR has surfaced as a practical alternative to randomized controlled trials (RCTs) in scenarios where implementation costs pose constraints and feasibility is restricted.[12] To investigate the causal relationship between relevant antibodies produced by H. pylori infection and skin disorders, we conducted a comprehensive MR analysis at the genetic level, applied UVMR and MVMR methods to to provide novel insights for the clinical treatment and alleviation of skin disorders.

## 2 Materials and Methods

### 2.1 Study Design

To ensure the credibility of our MR analysis, we used instrumental variables (IVs) adhering to three fundamental assumptions: (i) IVs should have a strong correlation with H. pylori antibody levels in exposure factors, explaining a minimum of 1.5% variation, with an F statistic surpassing 10 to mitigate bias from weak instruments; (ii) IVs should lack associations with confounding factors; and (iii) IVs should exclusively influence skin disorders through H. pylori antibody levels without genetic pleiotropy.[13,14]

Initially, genetic variants for seven antibodies specific to H. pylori were obtained from IEU OpenGWAS database. Subsequently, aggregated data on five skin disorders were derived from several GWAS datasets. We evaluated the causal link between H. pylori antibody levels and skin disorders through UVMR, MVMR, and various sensitivity analyses. Figure 1 illustrates the flowchart of MR performed in this study. As the data were pre-existing, no additional ethical approval was deemed necessary.

**Figure 1.**
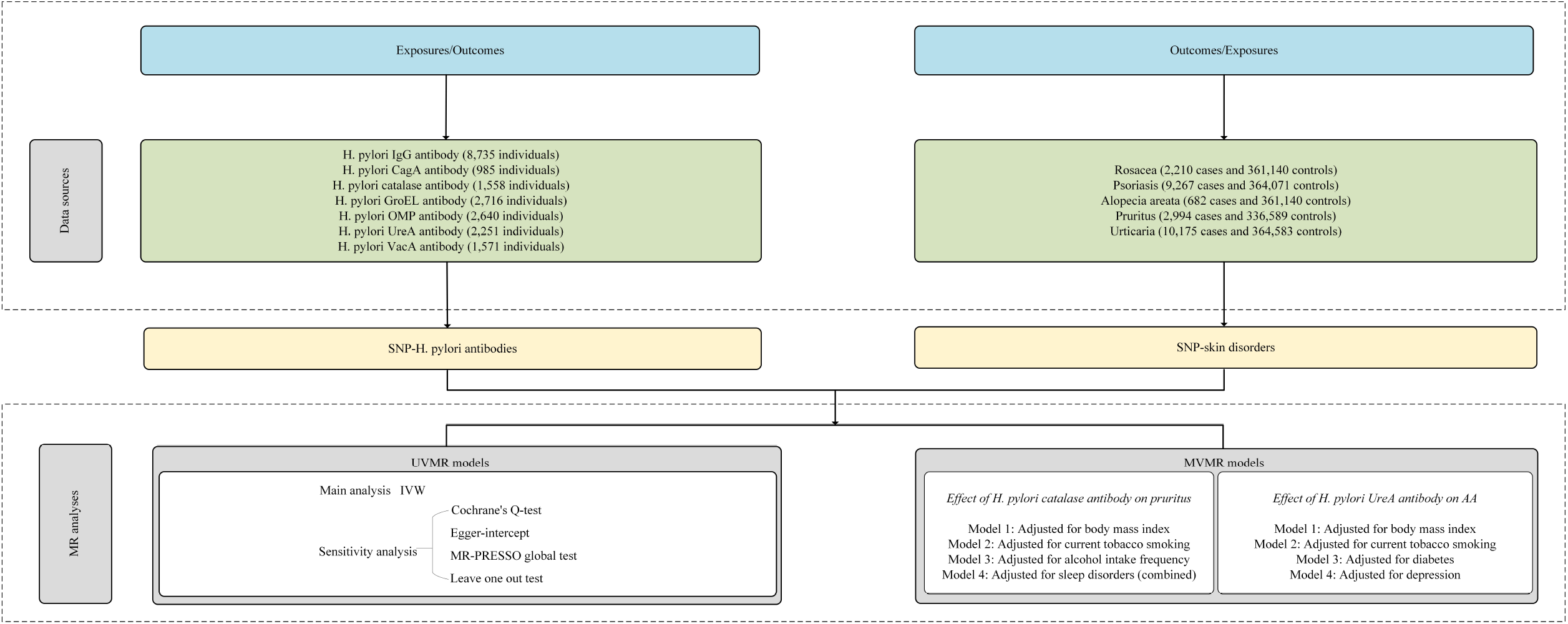
Flowchart of Mendelian randomization analyses conducted in this study. MR, Mendelian randomization. UVMR, Univariate Mendelian randomization. MVMR, Multivariate Mendelian randomization. IVW, Inverse variance weighted. SNPs, Single nucleotide polymorphisms. H. pylori, H. pylori. AA, alopecia areata. BMI, Body Mass Index.

### 2.2 Data sources

#### 2.2.1 Data for exposures

We obtained genome-wide association study (GWAS) data for seven distinct antibodies targeting H. pylori-specific proteins from the IEU OpenGWAS database (https://gwas.mrcieu.ac.uk/datasets/). These antibodies include anti-H. pylori IgG, chaperonin GroEL (GroEL), outer membrane protein (OMP), urease subunit-A (UreA), CagA, vacuolating cytotoxin-A (VacA), and catalase. IVs for exposure factors were derived from large-scale GWAS involving 985 to 8,735 participants of European descent, respectively. (Supplement Table S1)

#### 2.2.2 Data for outcomes

IVs for outcome factors such as rosacea, psoriasis, AA, pruritus, and urticaria were obtained from Finngen studies (https://r9.finngen.fi/) with 682 to 10,175 cases and 361,140 to 364,583 controls of European ancestry, respectively.[15] Notably, there was no sample overlap between the GWAS conducted for exposure and outcome. (Supplement Table S1)

#### 2.2.3 Data for confounders

We obtained genetic associations for BMI, current tobacco smoking, alcohol intake frequency, and sleep disorders (combined) from the IEU OpenGWAS database, while IVs for diabetes and depression were obtained from GWAS Catalog. (Supplement Table S1)

### 2.3 Selection of instrumental variables (IVs)

To investigate the causal relationship between exposure and outcome, we utilized single nucleotide polymorphisms (SNPs) as IVs. These selected SNPs needed to meet three MR assumptions. To secure an adequate number and statistical power of IVs, we established a p-value threshold for IVs at 5E-06 and organized them based on a linkage disequilibrium structure (r2 < 0.001, kb > 10,000 kb). A prerequisite for the F-statistic was that it be greater than 10; otherwise, it was deemed a weak IV and excluded. To ensure unbiased results, we implemented MR-Steiger filtering, removing SNPs highly correlated with the outcomes.[16,17] In the absence of SNPs in the final dataset, we employed TwoSampleMR to identify these missing SNPs and recognized alternative SNPs in high linkage disequilibrium with the primary SNPs (using a threshold of r^2^ > 0.8). It is crucial to highlight that SNPs must consistently be associated with the same alleles to accurately assess their impact on exposure and outcomes.

### 2.4 Statistical analyses

#### 2.4.1 UVMR analysis

Initially, we used the inverse variance weighted (IVW) method as the primary analytical approach to assess the causal connection between seven distinct antibodies targeting H. pylori-specific proteins and skin disorders. IVW utilizes a meta-analysis technique by combining the Wald estimates of each SNP, yielding an overall estimate of the exposure’s effect on the outcome. Under the assumption that all SNPs are valid instruments, IVW can offer the most precise estimation of the causal effect. Additionally, for enhanced reliability, we conducted supplementary analyses using the Weighted Median (WM) and MR-Egger methods.MR-Egger regression analysis showed the robustness to invalid instruments, and addressed the horizontal pleiotropy by introducing a parameter that accounts for potential bias.

The WM method collected up to 50% of the analysis information from the genetic variation of the inherent to zero instrumental variables.[18] The consistent estimation of the causal effect by three methods can enhance the causal inference.

To assess potential horizontal pleiotropy and heterogeneity that could substantially impact MR estimates, we conducted sensitivity analyses to enhance result credibility. (i) Cochran’s Q test was used to identify heterogeneity induced by horizontal pleiotropy and other biases. A P-value below 0.05 from the Cochran’s Q test suggested the presence of heterogeneity. In such cases, the random-effects model of the IVW method was employed to address heterogeneity.[19] (ii) MR-Egger intercept was used to gauge the extent of horizontal pleiotropy, with a p-value below 0.05 indicating its presence.[20] (iii) MR-PRESSO method was used to identify and correct outliers and horizontal pleiotropy.[21] (iv) Leave-one-out analysis assessed whether excluding any single SNP significantly affected overall MR estimates.[20]

#### 2.4.2 MVMR analysis

The MVMR analysis aimed to analyze the potential impact of confounding factors such as BMI, current tobacco smoking, alcohol intake frequency, and sleep disorders (combined) on the association between H. pylori catalase antibody levels and AA, as well as the potential impact of confounding factors such as BMI, current tobacco smoking, diabetes, and depression on the association between H. pylori UreA antibody levels and pruritus.

TwoSampleMR (version 0.5.7) and MRPRESSO packages (version 1.0.0) in R Software 4.3.1 (https://www.R-project.org) were used to perform the analysis.[18]

## 3 Results

### 3.1 IVs for MR

In the forward MR analysis, we obtained 3-13 SNPs as our IVs for the subsequent analysis (P < 5E-06, r2 < 0.001, kb > 10,000) based on the previously described stringent selection criteria, with explained variances ranging from 0.03% to 22.32%. SNPs that were directly associated with the outcome were excluded. F-statistic values were carefully considered and ranged from 21.11 to 28.17; no weak instrumental variables were present. (Supplement Table S2, S3)

### 3.2 UVMR analysis findings

Since IVW can provide the most accurate causal effect estimation, we based our assessment of significant causal relationships mainly on the IVW p-value. (Figure 2) In the UVMR, we identified a positive causal relationship between H. pylori catalase antibody levels and AA (IVW: OR = 1.32, 95% CI = 1.07 to 1.61, P = 0.008; WM: OR = 1.36, 95% CI = 1.03 to 1.79, P = 0.028). This analysis uncovered a 32% elevation in the risk of AA for every standard deviation (SD) unit increase in genetically predicted H. pylori catalase antibody levels; H. pylori UreA antibody levels and pruritus also had a positive causal relationship (IVW: OR = 1.17, 95% CI = 1.05 to 1.31, P = 0.006). Additionally, it substantiated a 17% rise in the risk of pruritus for every SD unit increase in genetically predicted H. pylori UreA antibody levels. However, MR-Egger and weighted mode analyses did not yield significant results (P > 0.05). (Supplement Table S2) After FDR correction, the above positive relationship became statistically non-significant, and the p-values for both IVW analyses were greater than 0.05. Indicating that there is a suggestive causal association. (Supplement Table S4) At the same time, the directions of IVW, MR-Egger, WM and Weighted mode were consistent (OR > 1), further supporting the suggestive causal association. Scatter plots depicting these two positive associations are presented in Figure 3. The relationship between H. pylori antibodies and other skin diseases, including rosacea, psoriasis, and urticaria is not significant (P > 0.05).

**Figure 2.**
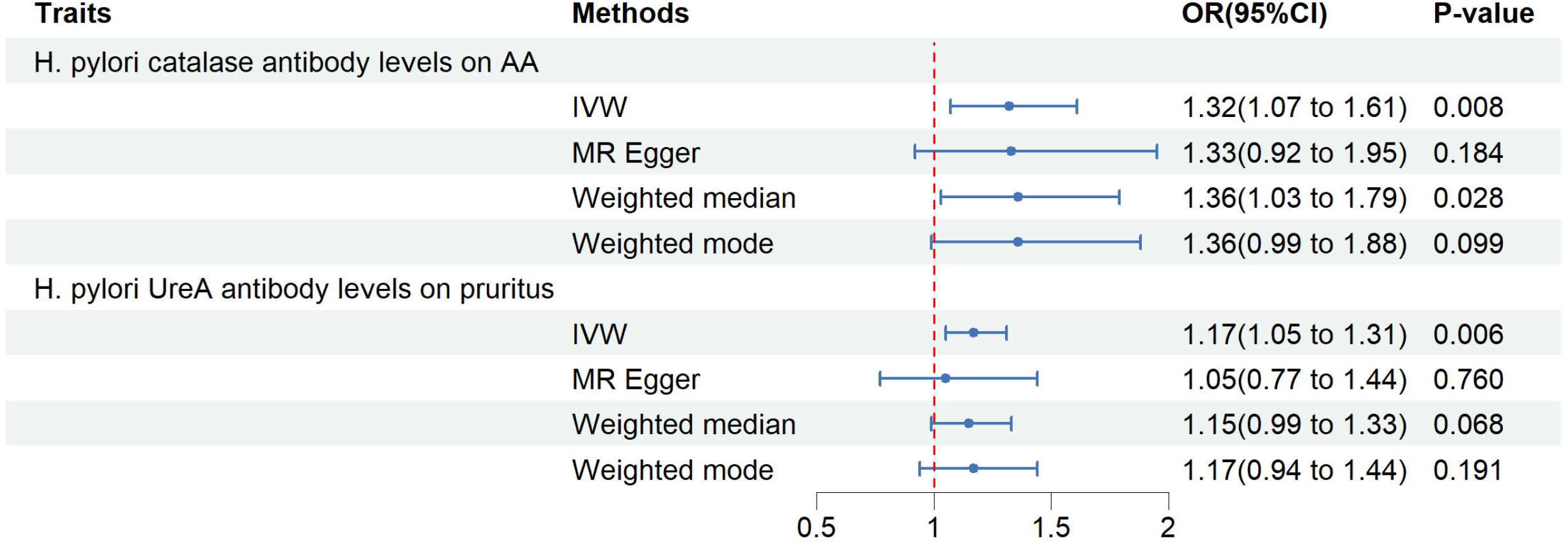
Associations of genetically predicted levels of H. pylori catalase antibody with the risk of AA and genetically predicted levels of H. pylori UreA antibody levels with the risk of pruritus. CI, confidence interval; OR, odds ratio.

**Figure 3.**
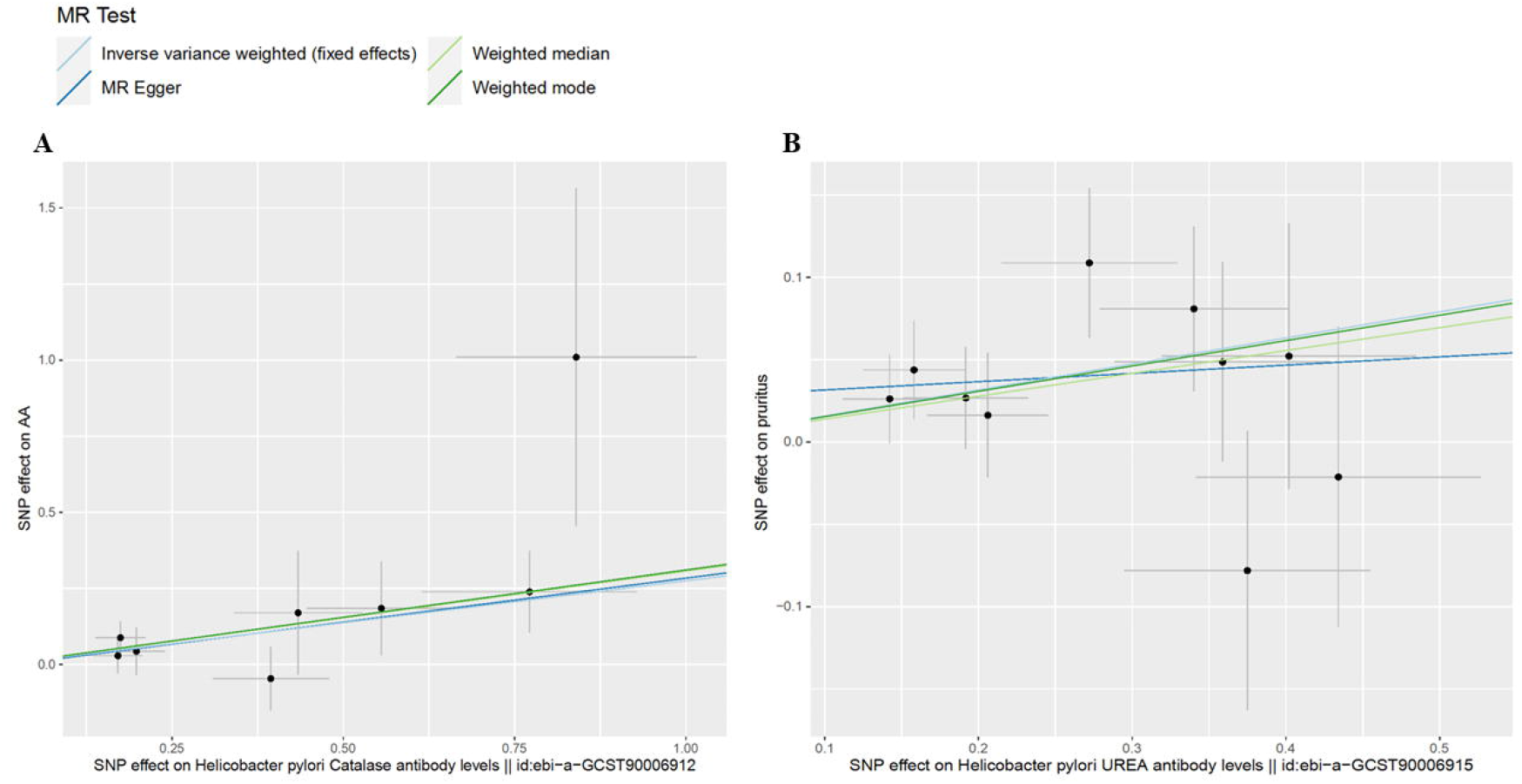
Scatter plots of two positive exposures (A) H. pylori catalase antibody (B) H. pylori UreA antibody levels.

### 3.3 Heterogeneity and sensitivity analysis

We conducted sensitivity analysis on the aforementioned significant findings. No confounding factors, including obesity, smoking, drinking, and diabetes, were identified in PhenoScanner (http://www.phenoscanner.medschl.cam.ac.uk/).[22-24] Cochran’s Q test revealed no heterogeneity among positive findings. The MR-Egger intercept suggested the absence of horizontal pleiotropy, while the MR-PRESSO test detected no outliers. Leave-one-out analysis indicated that eliminating any SNP would not impact IVW results. The MR-Steiger test confirmed an absence of highly associated SNPs with the outcome. (Supplement Table S5)

### 3.4 MVMR analysis findings

Using MVMR analysis, we found that the associations and significance of H. pylori catalase antibody levels with AA remained unchanged (P < 0.05) after adjusting for sleep disorders (combined). However, The estimated MR value of H. pylori catalase antibody levels lost statistical significance (P > 0.05) after adjusting for BMI, current tobacco smoking, and alcohol intake frequency, indicating that these factors are important as confounders. These factors may partly modulate the effect of H. pylori catalase antibody levels on AA. Similarly, the associations and significance of H. pylori UreA antibody levels with pruritus remained unchanged (P < 0.05) after adjusting for current tobacco smoking and depression. However, the estimated MR value of H. pylori UreA antibody levels lost statistical significance (P > 0.05) after adjusting for BMI and diabetes, suggesting that these factors are important as confounders. Hence, it is plausible that BMI and diabetes may, in part, mediate the impacts of H. pylori UreA antibody levels on pruritus. (Supplement Table S6)

## 4 Discussion

H. pylori, a gram-negative bacterium, is widely considered the causative agent behind peptic ulcer disease, gastric lymphoma, and gastric cancer. Its presence triggers robust white blood cell infiltration in the gastric submucosa, facilitated by pro-inflammatory cytokines. This pathogenic mechanism is recurrent in various other diseases. H. pylori seropositivity has been linked to diverse health conditions, including cardiovascular, respiratory, digestive, neurological, skin, and autoimmune diseases.[25,26] While the precise involvement of H. pylori in these diseases remains uncertain, the organism can be effectively eliminated through drug treatment. Research indicates that the resolution of various skin conditions, such as rosacea, chronic idiopathic urticaria, psoriasis, Behcet’s disease, and hereditary angioedema, can be achieved through the eradication treatment of H. pylori.[27]

The findings from our MR study revealed significant associations between certain H. pylori antibodies and AA as well as pruritus. However, no significant correlations were observed between H. pylori antibodies and rosacea, psoriasis, and urticaria in our investigation. Numerous prior studies have suggested a potential link between H. pylori infection and AA and pruritus. In the context of chronic skin diseases, persistent H. pylori infection may act as a triggering factor, exacerbating the conditions and complicating management. For instance, Belanghi E, Mansouri P, Agah S, et al., conducted a prospective study in the Iranian population involving 81 AA patients and 81 age and gender-matched healthy volunteers. Their findings indicated a higher prevalence of H. pylori infection in AA patients compared to controls, establishing its potential role in the pathophysiology of AA.[28] Campuzano-Maya G reported a case involving a 43-year-old male exhibiting patchy AA alongside an H. pylori infection. Following successful H. pylori eradication, the patient demonstrated hair regrowth.[29] Pruritus is a major symptom of skin diseases, which can adversely affect the health and quality of life of patients, and may lead to disability in severe cases. However, its mechanism is not well understood.[30,31] Previous reports have suggested an association between H. pylori infection and chronic pruritus, including cases of prurigo nodularis, and several studies have shown that the symptoms of pruritus are alleviated or even eliminated after H. pylori eradication.[4,32,33]

The concept of the gut-skin axis, supported by numerous studies, suggests that the gut microbiome can impact skin diseases through intricate immune mechanisms.[34] When a substantial quantity of intestinal microorganisms translocate, it exacerbates endogenous infection in the body, leading to immune system dysfunction. Simultaneously, metabolites from Staphylococcus, Streptococcus, Candida albicans, and Pityrosporum are generated on the skin. These metabolites directly interact with T cells through the human leukocyte DR antigen, activating the T cells. Subsequently, these activated cells migrate to the epidermis and swiftly release inflammatory mediators and epidermal growth factors. This process induces alterations in skin keratinocytes, ultimately resulting in the formation of skin lesions.[35] This represents the primary pathogenesis of the gut-skin axis. Concurrently, the intestinal mucosal immunity, being the largest and most intricate component of the body’s immune system, significantly influences overall immune function.[36] The essence of impaired intestinal mucosal immune function is that foreign antigens cause intestinal bacterial disorder through the tight junction of damaged intestinal epithelial cells, and trigger the immune response process, thereby giving rise to a disruption in the intestinal mucosal immune pathway, comprising mechanical, microbial, and immune cell components.[35-40] Autoimmune skin diseases arise from an immune system dysfunction, causing a breakdown in tolerance to skin autoantigens. The extended interplay between bacterial and host immune mechanisms suggests that H. pylori may act as a potential infectious agent, serving as a catalyst for the initiation of autoimmunity.[5] Thus, the unequivocal association between H. pylori infection and skin diseases is evident in their shared pathological mechanisms.

The treatment of many skin conditions, including AA and pruritus, is often a daunting challenge. Because it involves various treatment methods and the treatment time is long. Studies have shown that eliminating H. pylori can help treat some skin diseases.[5,27] Moreover, the understanding of the intestinal mucosal immune mechanism involving H. pylori and skin diseases remains unclear. Present-day medical research is primarily confined to theoretical and foundational investigations, lacking extensive clinical studies. Building upon current research findings, the establishment of a “gut-skin” axis model aligns with the contemporary scientific approach to treating autoimmune skin diseases like AA, showcasing limitless developmental potential. The adoption of the “gut-skin” axis concept in dermatosis treatment holds profound guiding significance and serves as a valuable reference for drug development. In the future, it is poised to achieve breakthrough significance in the multifaceted, multidimensional, and interdisciplinary approaches to preventing, monitoring, and treating skin diseases. Additionally, it may contribute significantly to dietary adjustments, healthcare practices, and psychological interventions in daily life.

The MVMR findings confirm that BMI and diabetes may partially mediate the effect of H. pylori UreA antibody levels on pruritus. Current research generally agrees that itching may be one of the symptoms associated with diabetes. The main cause of itching appears to be poor diabetes control, followed by dry skin and diabetic polyneuropathy.[41] Similarly, BMI, smoking, and alcohol may be confounders of H. pylori catalase antibody levels and AA. A study from Taiwan shows that smoking increases the risk of developing AA, while drinking alcohol can reduce the risk of AA.[42] Lifestyle-related factors have some influence on the pathogenesis of AA, including environmental factors such as smoking, alcohol consumption, sleep, obesity, fatty acid and gluten intake.[43] It can be inferred that improving lifestyle and controlling other diseases have potential benefits for pruritus and AA.

The advantages of our research are as follows: First, traditional observational studies have difficulty in drawing valuable causal inferences due to the presence of confounding factors, reverse causality, and small sample size. In our study, we employed genetic variants as instrumental variables to deduce the causal relationship between exposure and outcome, thereby mitigating the biases mentioned earlier.[44] Second, our IVs are derived from newly published articles and GWAS database. Our maximum sample size is up to 374,758, allowing us to better determine genome-wide risk and causal outcomes. Finally, our findings have potential implications for H. pylori treatment and medical policy in dermatology. In clinical application, improving H. pylori related examination and active treatment are beneficial to the recovery of some skin diseases.

Nevertheless, our study does have certain limitations. Firstly, the study population comprised individuals of European descent, potentially restricting the applicability of the findings to a broader global context. Secondly, the limitations of the GWAS dataset necessitate further expansion and enhancement of the sample size for certain skin disease studies. In the future, validation of our conclusions should involve a replication cohort with a more extensive sample size. Finally, there is a deficiency in research exploring underlying biological mechanisms linking H. pylori infection to AA and pruritus.

## 5 Conclusion

Our current findings reveal a positive causal relationship between H. pylori and AA and pruritus, which provides new perspectives and strategies for the clinical treatment of AA and pruritus.

## Data availability statement

All data used in the current study are publicly available GWAS summary data.

## Acknowledgments

Special thanks to the IEU open GWAS project developed by The MRC Integrative Epidemiology Unit (IEU) at the University of Bristol. Thank them for extracting relevant GWAS summary-level data from published articles, UK Biobank. And we also want to acknowledge the participants and investigators of the FinnGen study.

## Ethics statement

Ethical review and approval were not required for this study on human participants as it complied with local legislation and institutional requirements. Written informed consent was not obtained as the study utilized publicly available aggregate GWAS data. The study was exempt from Ethical Review Authority approval as it used public, anonymized, and deidentified data.

## Table captions

**Table 1.**
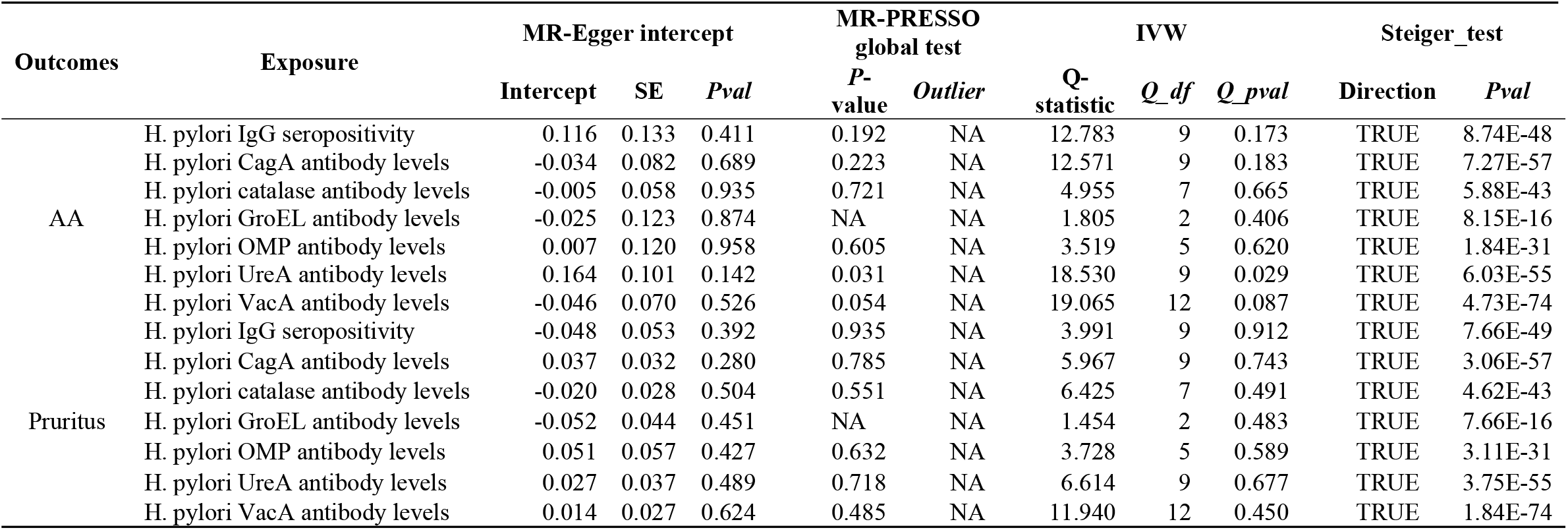
Sensitivity analysis of positive results. H. pylori: Helicobacter pylori; SE: standard error; IVW: inverse variance weighted; AA: alopecia area.

## Supplementary tables captions

**S1 Table**. Detailed information of data sources.

**S2 Table**. Analysis results of four methods of univariate Mendelian randomization (Effect of H. pylori antibodies on skin disorders, remove outliers).

**S3 Table**. SNPs information of H. pylori antibodies with skin disorders (p<5e-06, r2<0.001, kb>10000).

**S4 Table**. The positive results obtained from IVW method are corrected for FDR.

**S5 Table**. Summary of sensitivity results.

**S6 Table**. Summary of Multivariate Mendelian randomization results.

